# Relation of inflammatory marker trajectories with frailty and aging in a 20-year longitudinal study

**DOI:** 10.1101/2021.02.10.430670

**Authors:** Leonard Daniël Samson, Anne-Marie Buisman, José A. Ferreira, H. Susan J. Picavet, W. M. Monique Verschuren, A. Mieke H. Boots, Peter Engelfriet

## Abstract

Little is known about the development of low-grade inflammation with age and its relationship with the onset of frailty. In this exploratory study, we investigated 18 inflammatory markers measured in blood of 144 individuals aged 65-75 years at study endpoint, collected over 20 years at five-year intervals. IFNγ-induced markers and platelet activation markers changed in synchrony over time. Chronically elevated levels of IL-6-related markers, such as CRP and sIL-6R, were associated with frailty and becoming frail over time, poorer lung function, or less physical strength. Overweight was a possible driver of these associations. More and stronger associations were detected in women, such as between increasing sCD14 levels and frailty, indicating possible monocyte overactivation. Multivariate prediction of frailty showed low accuracy but confirmed the main results. In summary, we documented 20-year temporal changes of inflammatory markers in an aging population, and related these to clinically relevant health outcomes.

## 1 Introduction

Understanding the aging process is important for finding ways to prevent or delay age-related morbidity and thus to ensure good quality of life for an elderly population that is rapidly expanding worldwide (United Nations, 2017). One factor that may relate to ‘successful’ aging is an adequately functioning immune system, since an efficient immune response is essential to combat pathogens. Inflammatory processes can be out of balance in the elderly, leading to persistent low-grade inflammation (Baylis et al., 2013; Franceschi et al., 2017). It is thought that those with long-lasting low-grade inflammation have reduced responses to pathogens and carcinogenesis, and that they have more auto-inflammatory responses, and therefore being more prone to developing age-related diseases and to becoming frail (Baylis et al., 2013; Franceschi et al., 2017). One potential driver of chronic low-grade inflammation could be the amount of body fat, since adipocytes are thought to activate the immune system directly (Ghigliotti et al., 2014).

It is still largely unknown when and how low-grade inflammation develops during aging and how it is related to frailty. The few longitudinal studies performed on this subject showed that frail people often had low-grade inflammation for a long period of time (Gale et al., 2013; Puzianowska-Kuźnicka et al., 2016; Samson et al., 2019; Walker et al., 2018). In most studies, including our own (Samson et al., 2019), the presence of chronic low-grade inflammation was defined by the plasma concentrations of only one or two inflammatory markers, notably CRP and IL-6. While these are the markers most commonly used to investigate low-grade inflammation (Puzianowska-Kuźnicka et al., 2016), inflammation is a complex process in which many proteins are involved. Some studies already suggested that looking at a larger panel of inflammatory biomarkers, including a broader range of (chemotactic) cytokines, would improve the understanding of the relationship between low-grade inflammation and age-related diseases (Morrisette-Thomas et al., 2014). In addition, taking sex differences into account while studying the processes seems relevant because women generally have a higher frailty index score than men but do reach a higher age (Gordon et al., 2017).

In order to gain more insight into long-lasting low grade inflammation in relation to frailty and differences therein between men and women, we performed an exploratory study using data and blood samples from a selection of participants (n=144) of the longitudinal Doetinchem Cohort study. Blood samples and data were collected at 5-year intervals covering a period of approximately 20 years. We had several aims in our study. First, we wanted to explore the development of low-grade inflammation in an aging population, by investigating if and how concentrations of multiple inflammatory markers change with age. Secondly, we wanted to know if exposure to low-grade inflammation during a prolonged period of time relates to frailty or becoming frail, and to more specific aging-related clinical outcomes, such as decreased physical fitness (handgrip strength) and decline of lung function (spirometry). Lastly, we investigated if overweight is an important factor in these relationships.

## 2 Results

### 2.1 Study population characteristics

The study group consisted of 144 participants, (73 men and 71 women; Figure 1), with an average age of 68.3 years (min: 59.7, max: 73.5 years) at the endpoint of follow-up (Table 1). Average follow-up time was 19.3 years (min: 14.4, max: 20.9 years). Since blood samples were taken every five years, a maximum of five samples per individual was available for analysis, which was the case for most participants (n=107). Of the other study participants, either four samples (n=34) or three samples (n=3) were available. Highest concentrations were in the order of 10^6^ pg mL^-1^, for example of soluble CD14 (sCD14)(average endpoint level: 2.20 ∗10^6^ pg mL^-1^) and C-reactive protein (CRP) (1.22 ∗ 10^6^ pg mL^-1^) (Table S1), and lowest concentrations were around 1 pg mL^-1^, for example of interleukins such as IL-10 and IL-6 (average endpoint level: 0.65 and 2.31 pg mL^-1^, respectively). Concentrations of sIL-2R and IL-1β, were below detection limit in > 40% of the cases, and therefore these were excluded from analysis, leaving a total of 18 inflammatory markers for investigation (Table S1).

**Figure 1:**
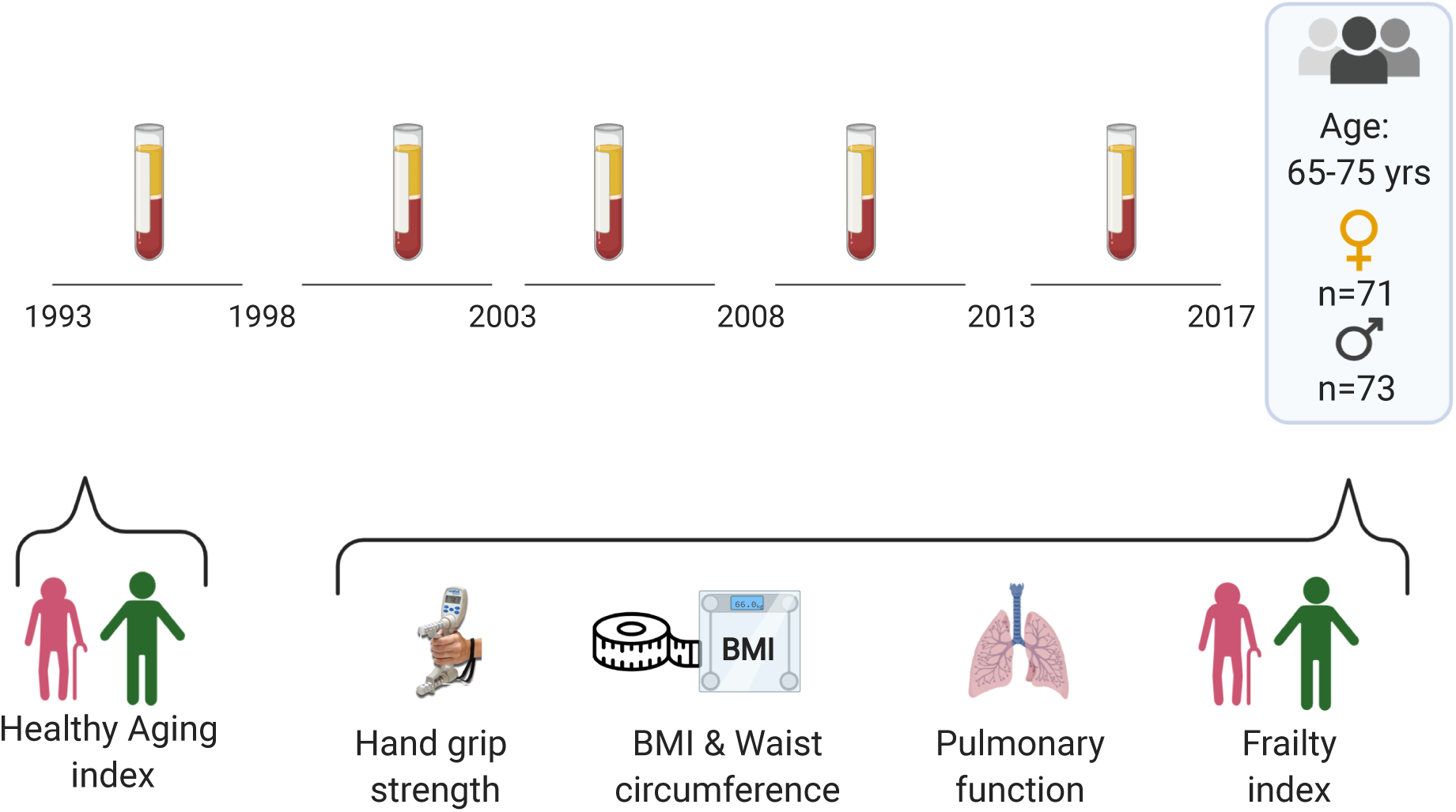
Timeline of the study. Participants of 65-75 years of age have been followed for more than 20 years, plasma samples were stored at 5-years intervals. At start, healthy aging index scores were determined. At last measurement, a frailty index on basis of 35 parameters including lung function and handgrip strength were determined which are used for separate analysis.

**Table 1:**
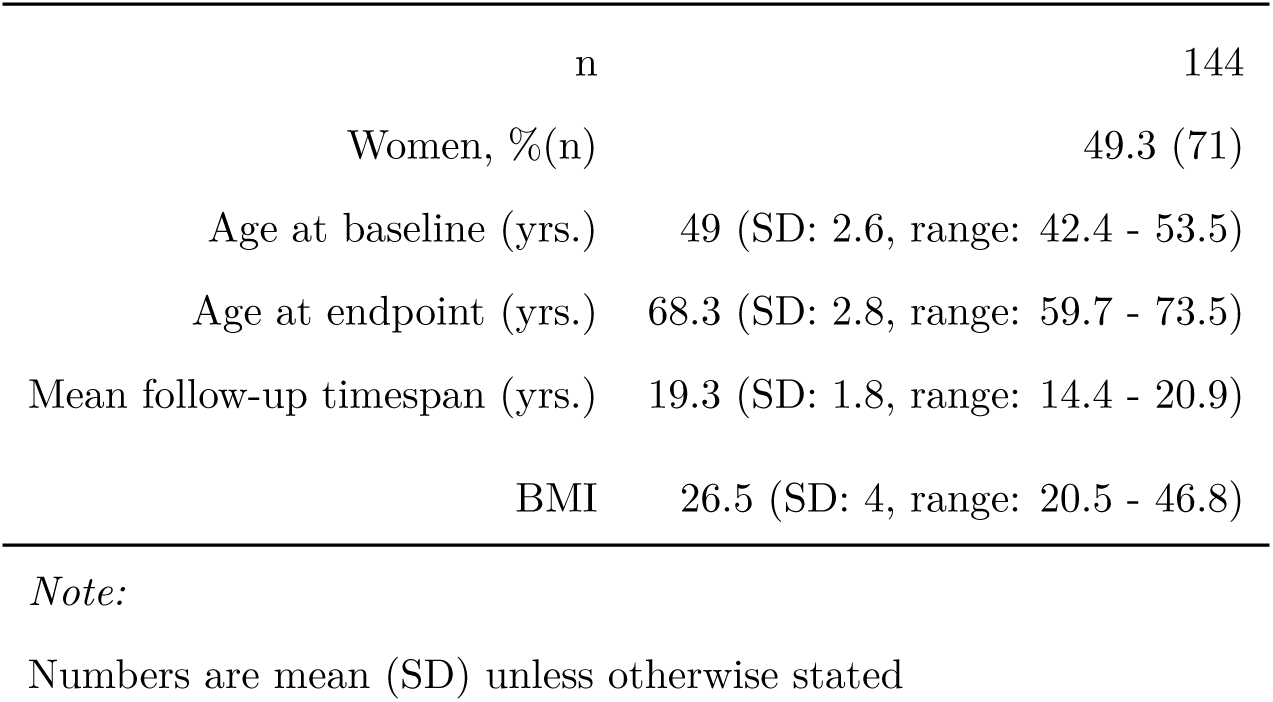
Baseline characteristics of the study population

### 2.2 Sex-specific changes in concentrations of inflammatory markers during aging

We calculated the average inflammatory marker levels over 20-year follow-up by means of the area under the concentration versus time curve (AUC) for each individual and each inflammatory marker separately. The AUC values differed between men and women for CCL5/RANTES, showing continuously higher levels in women (Figure S1). Several other dissimilarities between men and women became apparent when investigating the influence of age on the inflammatory marker levels. In men and women, an increase during aging was observed in the levels of CXCL10/IP-10, CXCL11/I-TAC and CCL27/C-TACK (Figure 2). In women, this increase was also seen for CRP, sIL-6R, CCL2/MCP-1, CCL11/Eotaxin, and sCD14, until age 60 approximately, after which age the trajectories of inflammatory markers in men and women appeared to become more similar (Figure 2). Indeed, women showed higher concentrations of CCL5/RANTES and BDNF, and lower concentrations of CCL11/Eotaxin and CCL2/MCP-1 at study baseline, while no differences were found between men and women in inflammatory marker levels at study endpoint (data not shown). The increase in inflammatory marker levels in women could be influenced by the major hormonal changes associated with menopause, since most women have their menopause before 60 years of age. Therefore, we compared the levels of inflammatory markers just before the menopause with those shortly after menopause in all women of whom these data were available at both timepoints (40 out of 70 women; Figure S2). Average self-reported age of menopause was 50.3 years (95% CI 39.7-60.9), and average time between measurement before and after menopause was 5.3 years (95% CI 2.6-8.1). Concentrations were higher after menopause for the inflammatory markers CCL2/MCP-1, CCL11/Eotaxin, sGP130, CCL27/C-TACK, and CXCL10/IP-10. Thus, our data suggest that inflammatory marker trajectories differ between men and women but become more similar with older age, which is possibly in part explained by hormonal changes in women during menopause.

**Figure 2:**
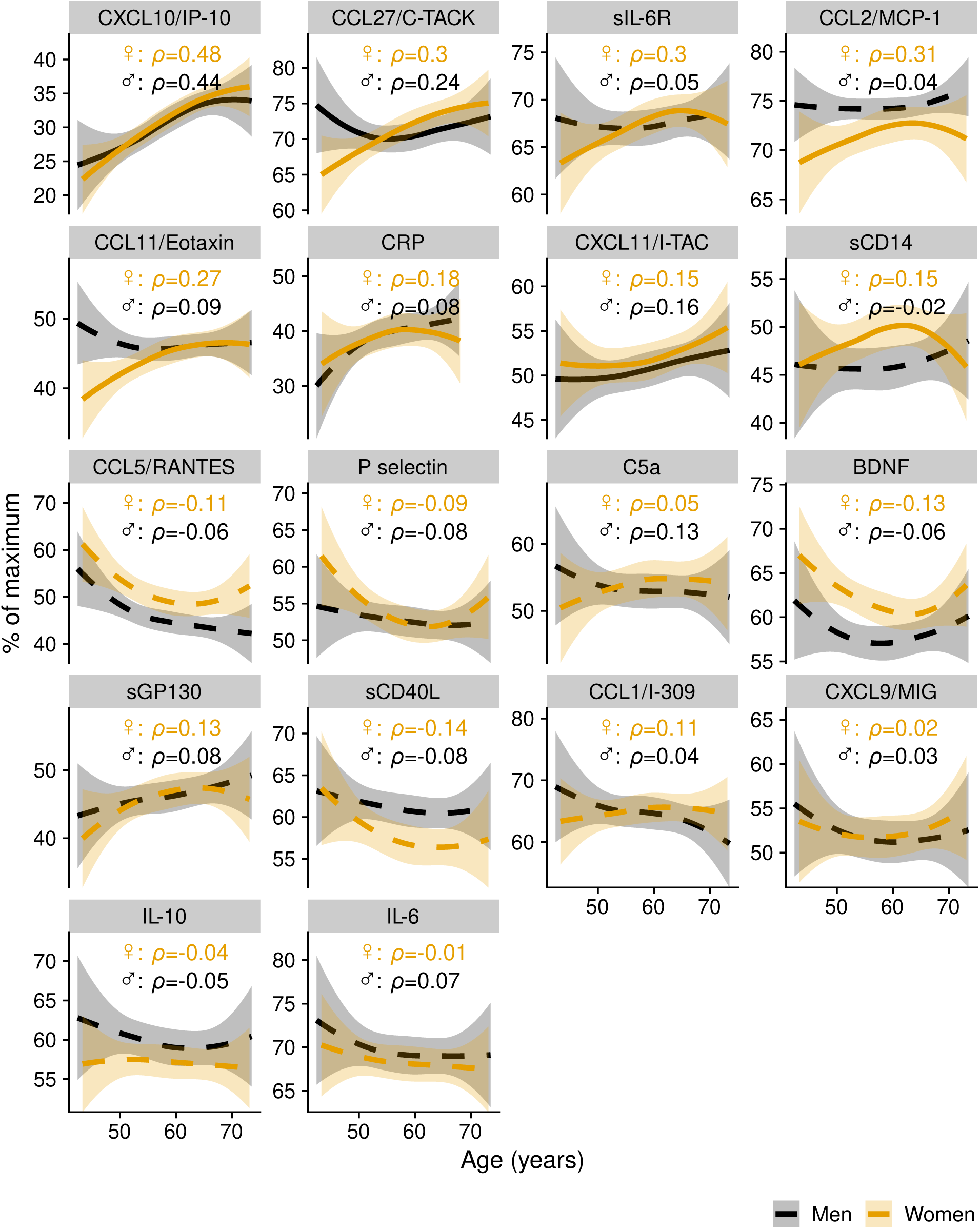
Trajectories of inflammatory markers related to age. The average trajectories are shown with 95% confidence interval estimated by local polynomial regression, separately for men (n=73) and women (n=71). Dashed line: no association found. Continuous line: association found between inflammatory marker trajectory and age. Y axis shows the % of maximum concentration per biomarker. Average concentrations per biomarker (pg mL^-1^) are indicated in Table S1.

### 2.3 Correlations between inflammatory markers at study endpoint

Testing of the correlations between inflammatory markers at endpoint revealed multiple associations (Figure 3A). All detected associations were positive, except for CCL27/C-TACK with P selectin, BDNF, sCD40L, and IL-10, and P selectin with sCD14. The strongest positive associations (*ρ*>0.50) were observed for IL-6 with IL-10 (*ρ*=0.72), IL-6 with sCD40L (*ρ*=0.58), IL-6 with CXCL9 (*ρ*=0.53), sCD40L with CXCL9 (*ρ*=0.50), and for BDNF with CCL5/RANTES (*ρ*=0.68). These associations were found both in men and in women but were stronger in women (Figure S3A,B). Other positive but weaker associations were seen between the structurally related IFNγ-inducible chemokines CXCL9, CXCL10/IP-10 and CXCL11/I-TAC (Figure 3A); again, these associations were stronger in women than in men (Figure S3A,B). Positive associations were found between BDNF and most of the CXCL- and CCL-chemokines (CCL1/I-309, CCL2/MCP-1, CCL5/RANTES, CCL11/Eotaxin, CXCL9, CXCL11/I-TAC) when men and women were analyzed together and stratified by sex (Figure 3A). When analyzing men and women separately, these were detected only in women (Figure S3A,B). Furthermore, weak correlations were found between several inflammatory markers involved in the IL-6 pathway, such as between sIL-6R and sGP130 (*ρ*=0.20), and between sGP130 and IL-6 itself (*ρ*=0.12) (Figure 3A). A weak association was found between higher levels of CRP and higher levels of sIL-6R (*ρ*=0.18) as well as with higher levels of CCL11/Eotaxin (*ρ*=0.20) (Figure 3A). When analyzed separately by sex, these relationships were detected only in women (Figure S3A,B).

**Figure 3:**
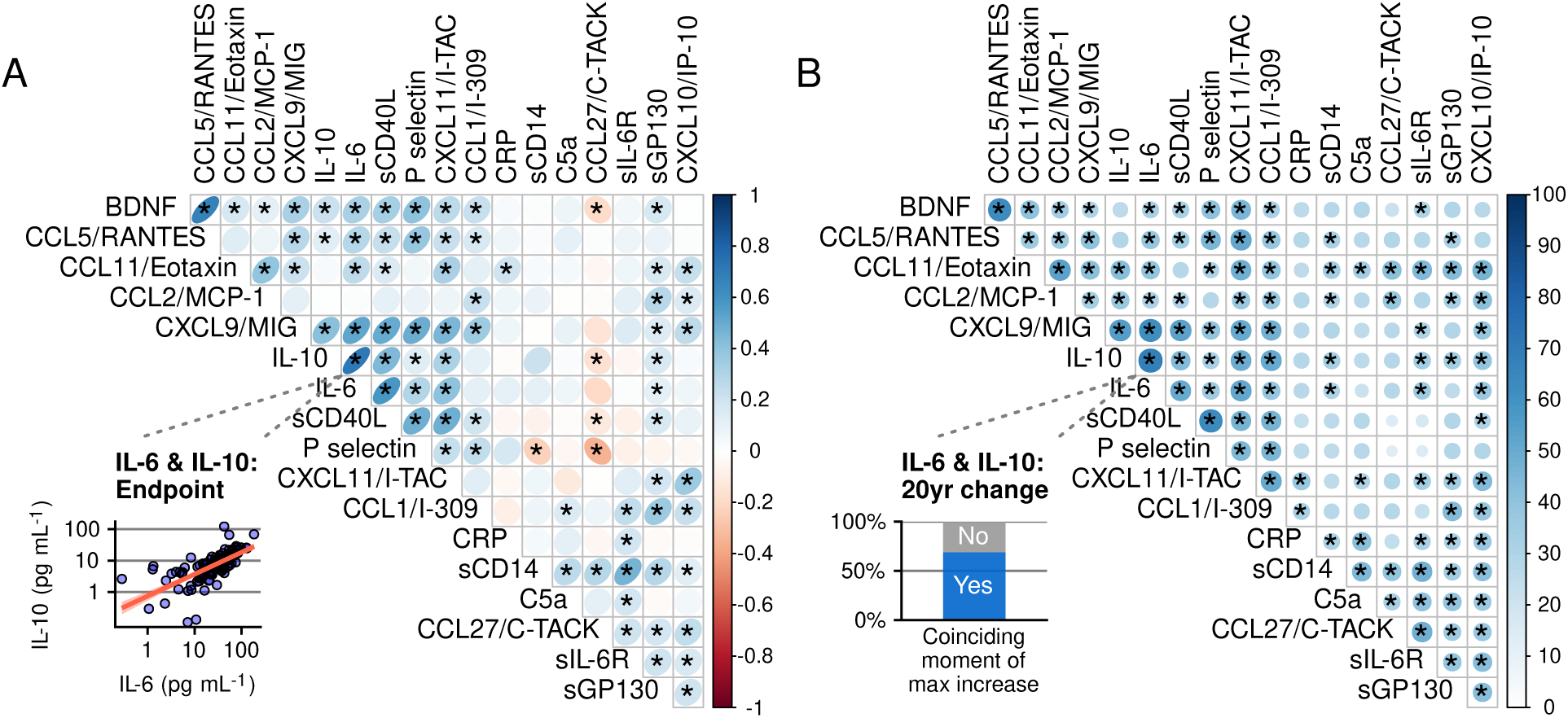
Relationships between inflammatory markers shown as A) correlation between pairs of inflammatory markers at study endpoint and B) similarity between pairs of inflammatory marker trajectories during about 20years of follow-up. In (A) the direction and strength of the association in is visualized with an oval shape and a color gradient. The inset shows the correlation between one pair of biomarkers (IL-6 and IL-10) as an example of the analysis. In (B) the blue gradient color and the size of the circles shows the percentage of participants of which a pair of biomarkers had the highest increase in concentration at the same moment in 20 years of follow-up. This was seen in 69% of the participants for the pair of biomarkers IL-6 and IL-10 (see inset). *= an association between two inflammatory markers, with false discovery rate being set at a maximum of 15%. n=144

### 2.4 Similarities between inflammatory markers in terms of temporal changes over 20 years

Since there is high variability between individuals in concentrations of inflammatory markers, a cross-sectional analysis is somewhat restricted. Therefore, we looked for synchronous changes of biomarkers, meaning that when biomarkers change together this is probably caused by an underlying biological process (in response to a stimulus, or under homeostasis) inducing these changes. To do so, we determined in which of the four possible time intervals in the 20 years of follow-up time a marker had its greatest increase in concentration. Then, we tested if this moment was the same for every pair of biomarkers in the majority of the participants. In both men and women the most distinct synchronous changes were again found with IL-6 and IL-10 (Figure 3B), and with markers of the innate immune system, such as those of platelet activation (sCD40L and P-selectin) and those related to chemotaxis and granulocyte activation (CCL11/Eotaxin and CCL5/RANTES) (Figure 3B). The concentrations of structurally related chemokines CXCL9, CXCL11/I-TAC and, to a lesser extent, CXCL10/IP-10 also tended to change in unison over time in both men and women. In addition, CXCL9 and 11 were related to multiple other inflammatory markers, such as IL-10 and IL-6, but also RANTES/CCL5, P selectin, and sCD40L, indicating the sensitivity of inflammatory marker levels for ifN-pathway activation in aged men and women.

In this longitudinal analysis inflammatory marker profiles of men and women appear somewhat more similar than in the cross-sectional analysis. Thus, while RANTES/CCL5 correlated with more markers in women than in men looking at the marker levels at study endpoint (Figure S3A,B), this appeared to be less so when looking at synchronous changes in markers over time (Figure S3C,D). Furthermore, we did not find differences between men and women in the maximum levels of biomarkers over the entire 20-year time period, nor in the magnitude of the maximum increase in inflammatory marker levels (data not shown). In summary, multiple inflammatory marker levels were correlated with each other, with IL-6 showing the strongest correlations with other markers, followed by markers related to platelet activation and the IFNγ-pathway activation. Correlations between inflammatory markers at endpoint were stronger and more abundant in women, but this tendency was less pronounced when comparing the temporal change in biomarker levels, with no strong evidence found for higher immune marker reactivity in women.

### 2.5 Relationships of inflammatory markers with BMI and waist circumference

We investigated the relationship between the inflammatory marker profile and two measures of body fat, namely body mass index (BMI) and waist circumference. The average BMI increased with age in both sexes over the 20 years of follow-up (men: *ρ*=0.24, women: *ρ*=0.37, Figure S4A). In women, the average BMI values and waist circumference during the follow-up were positively associated with the AUC of CRP (*ρ*=0.55) and sIL-6R (*ρ*=0.28) (Figure S4B and C), both related to the IL-6 pathway. In men, a larger waist circumference was positively associated with CRP levels, but associations between average BMI levels and inflammatory markers were not observed.

### 2.6 Levels of inflammatory markers related with frailty

Continuously elevated CRP trajectories were related to frailty at study endpoint (frailty index based on 35 deficits, Table S2) as was shown previously (Samson et al., 2019), and this association turned out to be one of the strongest when analyzed together in our inflammatory marker panel (correlation of study endpoint frailty with the area under the curve (AUC) of the CRP trajectory in men: *ρ*=0.44, in women: *ρ*=0.50; see Figure 4A&B). Associations of frailty with other inflammatory markers were only found in women, showing a higher AUC of sIL-6R (*ρ*=0.34) and sCD14 (*ρ*=0.33), but lower AUC of sCD40L (*ρ*=-0.29) in frailer women. BMI values were also higher on average at each time of measurement in frail men and women (Figure 4C). When associations of frailty with the inflammatory markers were adjusted for BMI, two correlations were found, namely a positive association of CRP values with frailty and an association that was not found without adjusting for BMI, namely a positive one of BDNF with frailty in men (*ρ*=0.35 and *ρ*=0.21, respectively). We further investigated whether age-related increases or decreases of inflammatory markers over the past 20 years were specifically related to frailty at study endpoint. The age-related increases in sCD14 and sIL-6R levels seen in women (Figure 3) turned out to be related to frailty at study endpoint (Figure 4D, *ρ*=0.42 and *ρ*=0.35, respectively). In addition, while no age-related increase in C5a levels were seen, C5a levels were seen to increase with age with higher frailty index scores in women (Figure 4D, *ρ*=0.30). These results were also found after adjusting for both BMI and for baseline marker concentrations.

**Figure 4:**
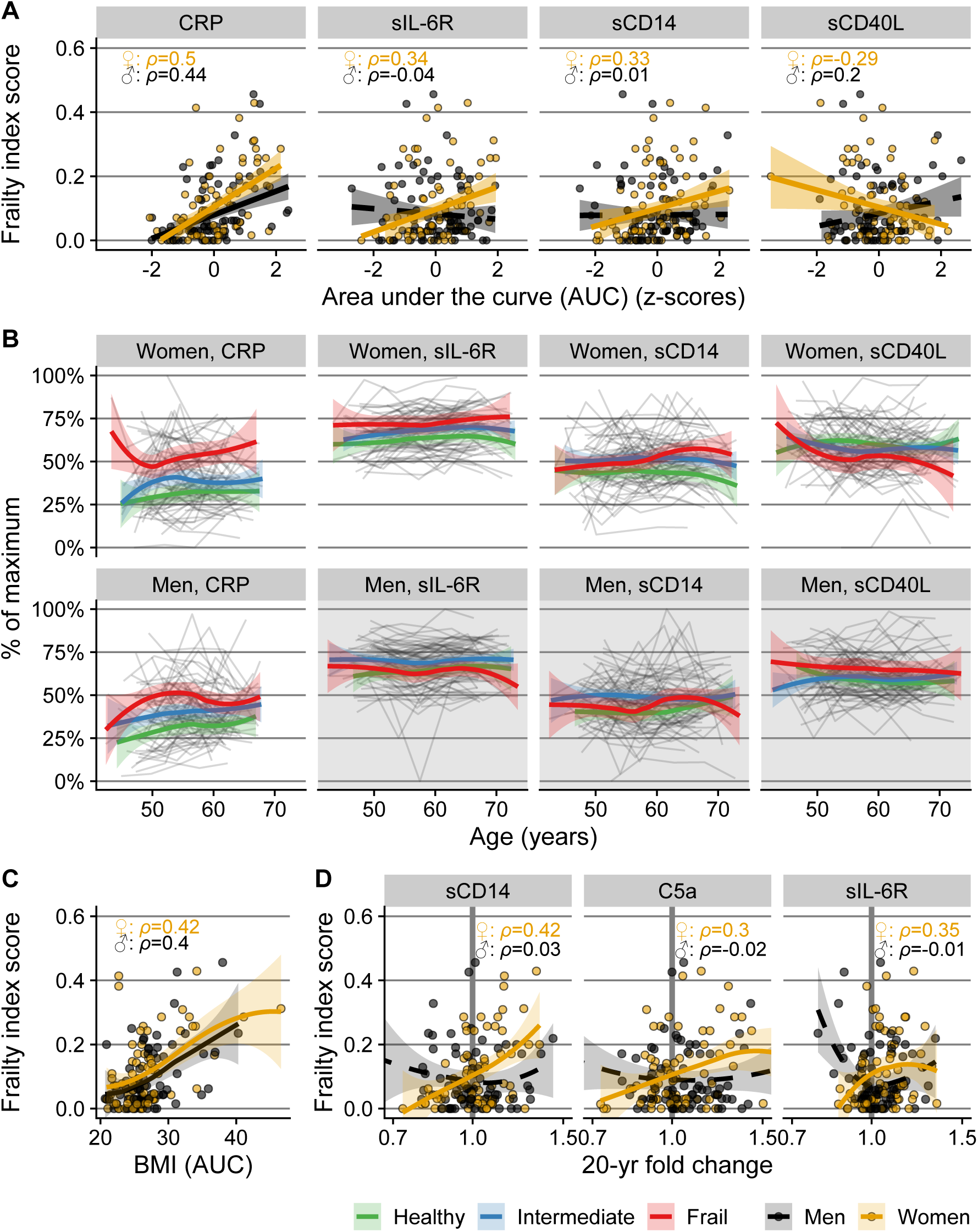
Relation between frailty at endpoint and inflammatory marker levels over time in men (n=73) and women (n=71). To capture the cumulative ‘exposure,’ the inflammatory markers in (A) are expressed as area under the inflammatory marker concentration curve versus time (AUC), standardized to take into account different follow-up periods, and transformed into z-scores for visualization and better comparison between inflammatory markers. (B) shows the trajectories of individuals (grey lines) and the local polynomial regression lines with 95% confidence intervals per frailty category (bold colored lines) based on frailty index score at study endpoint (Concentrations are scaled to % of maximum concentration per marker). This complements the analysis in (A) and visualizes in more detail whether marker levels are elevated chronically. The frailty categories in (B) were used for illustration purposes; for the statistical analyses, the continuous frailty index score was used. Inflammatory markers were displayed when their AUC showed an association with the frailty index score at endpoint in at least one of the sexes. Plot area backgrounds in (B) are grey if no association was found. (C) Body mass index (BMI) over time, expressed as AUC values. (D) 20-year fold change in inflammatory markers, all over 20 years. A vertical reference line of no increase (fold change of 1.0) is shown in bold grey.

### 2.7 Prediction of frailty with multiple inflammatory protein trajectories

Since multiple associations were found between inflammatory protein trajectories and frailty, we further investigated this relationship by studying how well the AUCs of all the biomarkers together could predict the frailty index score, using a random forest algorithm with BMI and age at last measurement also included in the model. The predictive accuracy was 12% in men and 10% in women. With this low predictive accuracy, only the top results of the algorithm could be interpreted (Figure 5).These top results confirmed findings of the association study, with BMI and CRP being the most important predictors with comparable variable importance in men and women, and sCD14 closely following albeit only in women.

**Figure 5:**
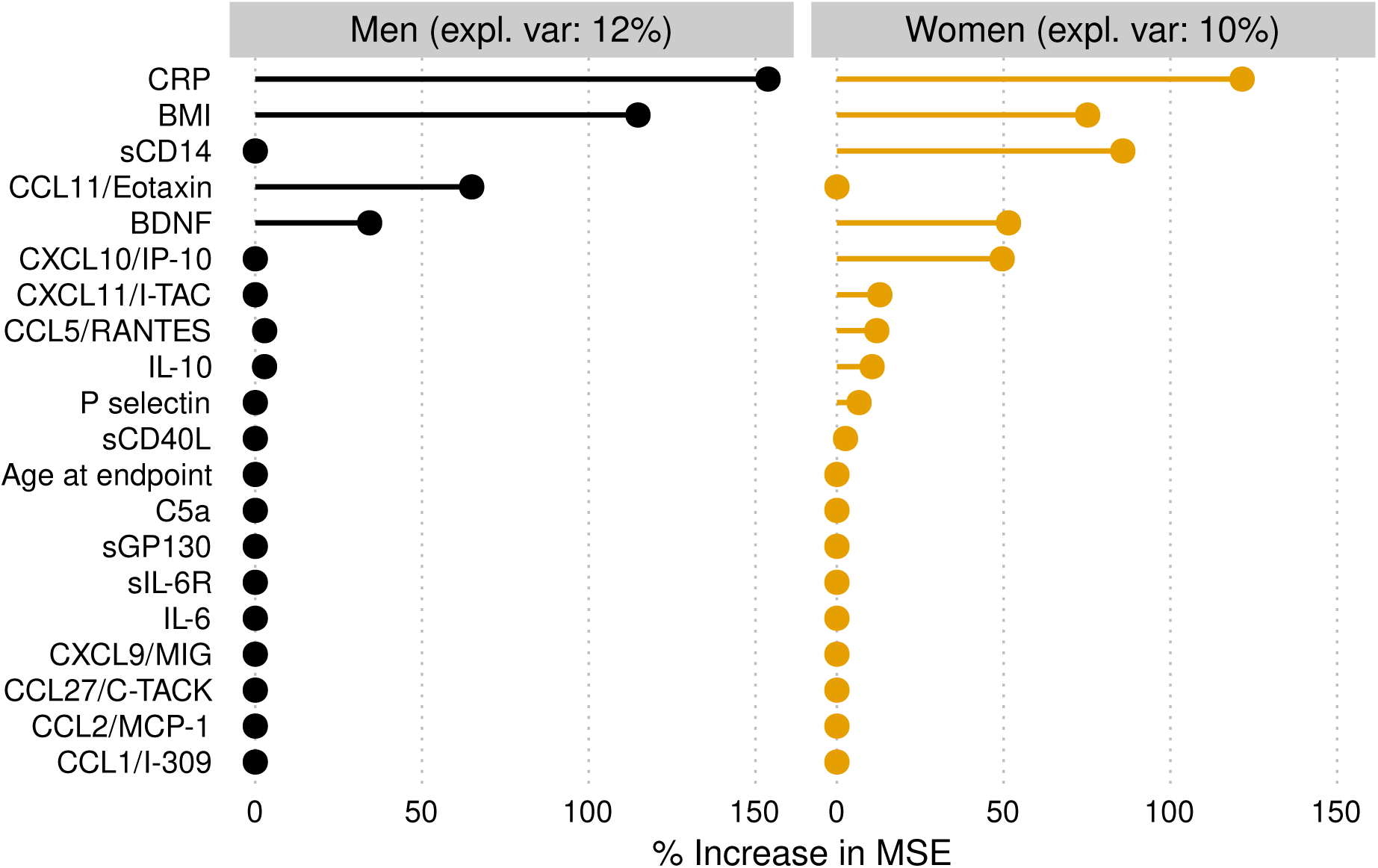
Variable importance of the inflammatory marker concentrations over the past 20 years (area under the curve) per individual in predicting frailty at study endpoint using a random forest algorithm. Age at study endpoint was included in the model. Expl. Var: explained variance. % increase in MSE: percentage increase in mean squared error of the prediction of frailty after the variable is replaced by random noise. A higher value thus means that the variable is more important in predicting frailty.

### 2.8 Trajectories of inflammatory markers related to an increase in frailty

Next, the relation was studied between the development of low-grade inflammation and the onset of frailty. This was done in two different ways. First, we investigated whether chronic low-grade inflammation was related to an increase in frailty over a period of five years, i.e. between measurement rounds 5 and 6. While the AUCs of several inflammatory markers were found to be related to frailty, no associations were observed between them and the increase in frailty over the five-year period (Supplementary correlation tables, tab “Frailty_change”). Secondly, to investigate the risk of becoming frail prospectively over a longer period of time, we selected a subgroup of individuals who were ‘healthy’ and thus were likely to have a low, favorable frailty index score at the start of our study; healthy at study baseline was defined according to an alternative health index that could be assessed at study baseline (Healthy Ageing Index score, score of 9 or 10 out of 10, higher score being more favorable; 31 women and 27 men; Figure 6A) (Dieteren et al., 2020). Then we investigated within these selected ‘healthy’ participants whether those with continuously higher (or lower) levels of inflammatory markers had a higher frailty index score at study endpoint, and thus became frailer over time. In this subgroup, we observed that in women frailty development was associated with higher AUCs of CRP and sIL6R (*ρ*=0.44 and 0.56 respectively, Figure 6B&C). In men, no associations with frailty were found. In the relationship between inflammatory markers and becoming frail body fat could be important, since BMI was continuously higher in participants who became frail (Figure 6D). Indeed, after adjusting for BMI, the associations of the inflammatory markers with the chance of becoming frail in women were weaker and no longer passed the criteria for qualifying as a “detected association” (correlation coefficient of frailty and sIL-6R: *ρ*=0.49; of frailty and CRP: *ρ*=0.22). Thus, it is likely that BMI plays an important role in the association of chronic inflammation over time and frailty.

**Figure 6:**
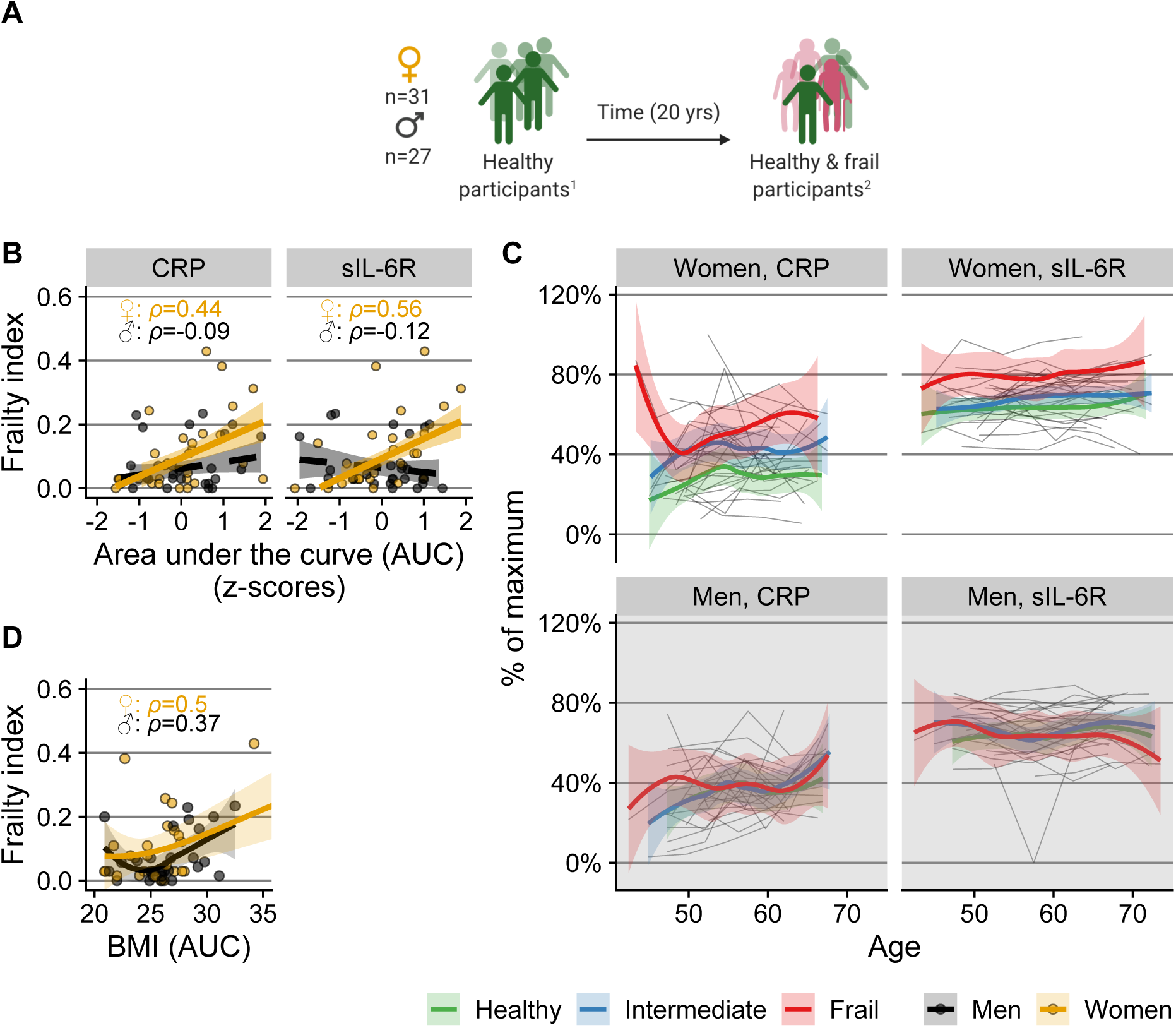
(A) Subgroup of participants that were ‘healthy’ at baseline. ^1^Health status at baseline is defined by a healthy aging index (healthy aging index score = 9 or 10 out of 10). ^2^Health status at endpoint is defined by a frailty index score. Relation of frailty at study endpoint with trajectories of CRP and sIL-6R (B,C) and with BMI (D) in the subgroup of people that were ‘healthy’ at study baseline. To capture the cumulative ‘exposure,’ trajectories of CRP and sIL-6R in (B) and of BMI in (D) are expressed as area under the concentration/level versus time curve per individual. Grey lines in (C) are individual trajectories. Bold colored lines in (B) are (robust) linear regression lines, and in (C) and (D) local polynomial regression lines with 95% confidence intervals. In (C), the color denotes the ‘frailty category’ at study endpoint. Frailty categories are used for visualization of the longitudinal trajectory; for the statistical analyses the continuous frailty index score was used. Concentrations on y-axis in (C) are scaled to % of maximum concentration per marker.

### 2.9 Trajectories of inflammatory markers related to lung function

As lung function is an important determinant of health known to decline with age, and for which clinically accepted quantitative measures have been defined, we investigated the relation between inflammatory marker trajectories over the past 20 years and spirometric measurements at endpoint. In women, pulmonary function, in terms of Forced Expiratory Volume in one second (FEV1), tended to be lower when trajectories of CRP were higher (correlation of the AUC of CRP with FEV1, *ρ*=-0.48),but higher with elevated trajectories of CXCL11/I-TAC (*ρ*=0.33), and sCD40L (*ρ*=0.33) (Figure 7A). In men, no associations were found between the inflammatory marker trajectories and the FEV1, although the patterns of CRP were more or less similar to those in women. A different pulmonary function measure, Forced Vital Capacity (FVC), was found to be negatively related to CRP concentrations in women (*ρ*=-0.48) adding to the notion that pulmonary function is affected by chronic inflammation. No relationships were found between the inflammatory marker trajectories and the FEV1/FVC ratio.

**Figure 7:**
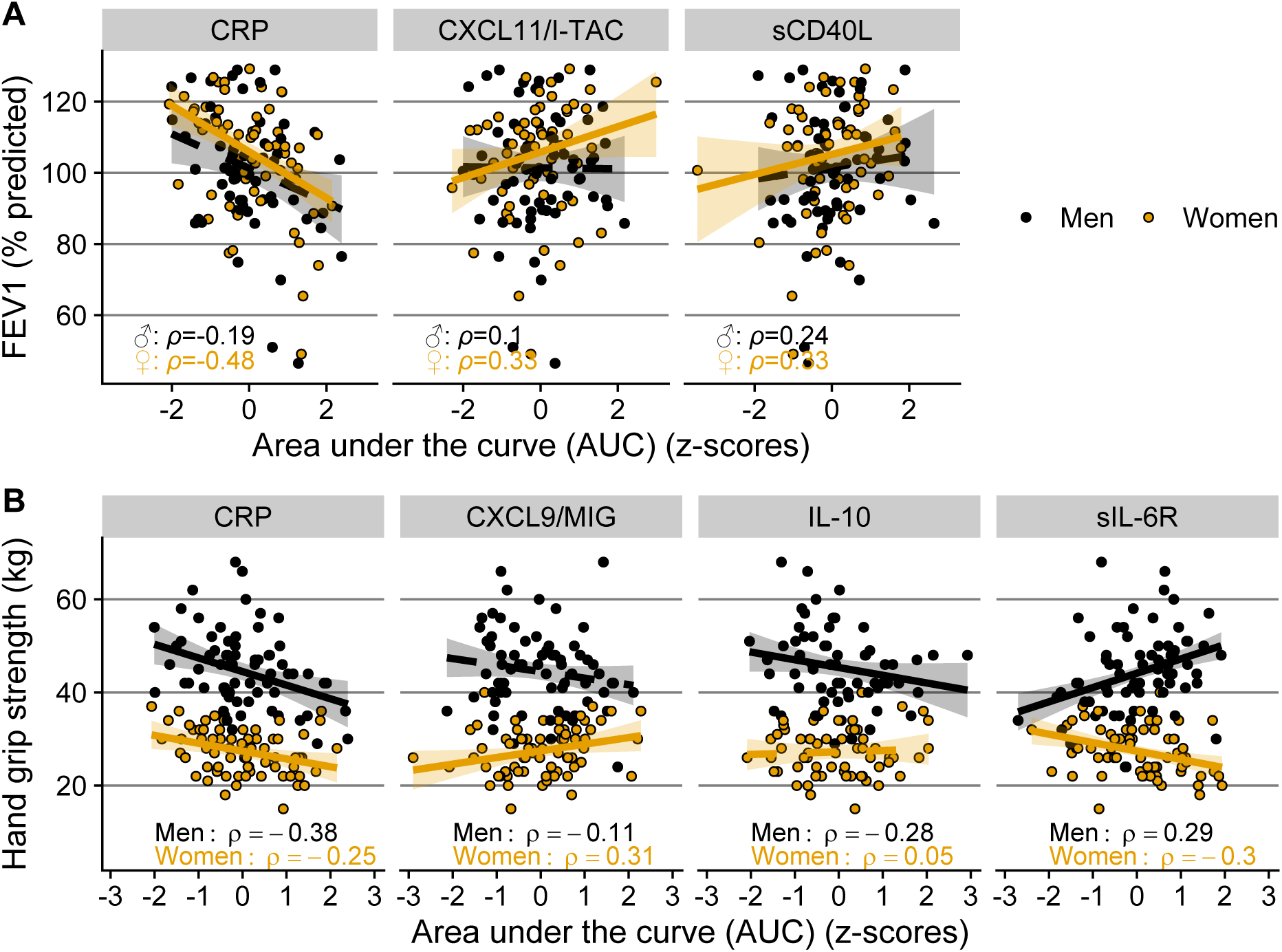
Inflammatory marker trajectories in men and women of those that showed an association with physiological clinical parameters of aging at study endpoint; (A) lung function assessed by forced expiratory volume in one second (FEV1), and (B) handgrip strength measured as an approximation of physical strength. As a measure of cumulative ‘exposure,’ inflammatory marker levels were assessed as area under the inflammatory marker concentration versus time curve per individual (AUC) during the 20-year follow-up. AUC values were transformed to z-scores for visualization. % predicted: FEV1 value compared to expected reference values from the Global Lung Initiative (Quanjer et al., 2012) specific for age, length, and sex. An association is indicated by a continuous trendline, otherwise a dashed trendline is shown.

### 2.10 Trajectories of inflammatory markers related to handgrip strength

Another important determinant of health that declines with advancing age is muscle strength, which can be represented approximately by handgrip strength. Handgrip strength, as measured with a dynamometer, was related to several inflammatory marker trajectories in men and in women (Figure 7B). In both sexes we found a negative association between the AUCs of CRP trajectories and handgrip strength, but only in men did this association remain after adjusting for BMI. In men, a negative association was found between IL-10 trajectories and handgrip strength (*ρ*=-0.28), which remained after adjusting for BMI. Contrasting correlations were found between sIL-6R and handgrip strength in men and women, with men showing a positive correlation (*ρ*=0.29) and women a negative one (*ρ*=-0.30). After adjusting for BMI, however, these associations with sIL-6R were lost. Women also showed a positive correlation of handgrip strength with CXCL9 (*ρ*=0.30) which was lost after adjusting for BMI. Taken together, chronically elevated levels of CRP and IL-10 are associated with lower hand grip strength in men.

## 3 Discussion

In this unique longitudinal study with 20 years of follow-up data, we quantified chronic low-grade inflammation in an aging population using multiple inflammatory markers and showed that changes of markers of the IL-6 pathway, of platelet activation and of monocyte activation are associated to clinically relevant health outcomes at older age, such as frailty. BMI and waist circumference turned out to be important factors in this process, since they are related to both frailty and markers of the IL-6 pathway. Associations were stronger and more abundant in women, although inflammatory marker trajectories in men and women became more similar with age. Although the power to predict frailty using the inflammatory marker trajectories was weak, probably due to the heterogeneity in the study population and the small numbers of individuals, the results were in agreement with those of the association studies.

### 3.1 Age-related increase of inflammatory marker levels

We detected an increase in multiple biomarker levels with advancing age, which is in line with the reported chronic low-grade inflammation at older age (Baylis et al., 2013; Franceschi et al., 2006). Clear increases in ifN-related chemokines with age were seen in both men and women. Higher concentrations of inflammatory markers with advancing age were seen in both men and women in the IFNγ-induced and structurally related chemokines CXCL10/IP-10 and CXCL11/I-TAC. Elevated concentrations of these chemokines could be a sign of both innate and adaptive immune activation and could be clinically important, since they have been proposed as possible biomarkers for heart failure (Altara et al., 2015). CXCL10/IP-10 has previously been described as increasing with age in studies with shorter follow-up time (Hearps et al., 2012; Hsu et al., 2019). Also CCL27/C-TACK levels increased with age in men and women, possibly related to accumulating skin damage in elderly, since CCL27/C-TACK is a cytokine involved in T-cell-mediated homing to the skin (Homey et al., 2002; Richmond et al., 2019).

In women, other inflammatory markers were also found to increase with advancing age, including the IL-6 pathway related markers CRP and sIL-6R and two chemokines related to innate immune cell activation, namely CCL2/MCP-1 and CCL11/Eotaxin. These sex-specific increases all seemed to reach a plateau at around the age of 60 years. Differences in the immune profile between men and women could be due to hormonal differences, as stated previously (Bupp, 2015; Furman et al., 2014). This is in line with our data suggesting that hormonal shifts due to menopause can partially explain why concentrations of multiple inflammatory markers increase over time in women before the age of 60. Of note is that, while sIL-6R and CRP were found to increase with age in women, IL-6 levels were not. This may be explained by the molar excess of sIL-6R in the circulation, which naturally binds IL-6. Still, we expected to find an association between IL-6 levels and age, since IL-6 levels were found to be increased with advancing age in several previous studies (Ferrucci et al., 2005; Giuliani et al., 2001; Puzianowska-Kuźnicka et al., 2016), which is why it is thought to be a key component in chronic low-grade inflammation in the elderly. There are, however, other studies that also did not find a relationship between IL-6 and age (Beharka et al., 2001; Hsu et al., 2019; Van Epps et al., 2016), one of which claimed this was due to the limited age range of their participants (Van Epps et al., 2016).

### 3.2 Chronic low-grade inflammation related to frailty

In our study we investigated how chronic low-grade inflammation is related to frailty and to an increase in frailty. As expected, the association of higher CRP levels in frailer men and women that we found in our previous study (Samson et al., 2019) was detected in this subcohort again, even in a smaller selection of participants (n=144 instead of 289), and with a different analysis approach, i.e. by investigating the area under the curve of the inflammatory markers. Moreover, CRP turned out to be one of the inflammatory markers with the strongest association with frailty and with the chance of becoming frail over time, followed by sIL6R in women, indicating a role for the IL-6 pathway. Previous literature about immune marker alterations preceding the risk of becoming frail is limited, with one study showing associations of higher CRP and the risk of becoming frail (Gale et al., 2013). In another study, associations were reported between IL-6 and ‘incident frailty’ over 5 years. sIL-6R was not measured in that study (Hsu et al., 2019). Probably a major driver in this process is (over)weight, since the associations with frailty in our study were not detected anymore after adjustments for BMI. A plausible reason why BMI influences these results is that a higher BMI both increases the risk of becoming frail as well as raises the inflammatory biomarker levels (Ghigliotti et al., 2014). Excessive adipose tissue, especially visceral adipose tissue, is known to contribute to chronic inflammation by increased secretion of adipokines and inflammatory cytokines and by promoting accumulation of macrophages (Ellulu et al., 2017; Ghigliotti et al., 2014; Ouchi et al., 2011). It has been inferred that waist circumference is a better estimate of visceral pro-inflammatory body fat than BMI in the elderly (Crow et al., 2019). Indeed, we saw stronger associations of IL-6 pathway markers with waist circumference than with BMI in men, possibly because waist circumference is thought to give a better estimate of visceral pro-inflammatory body fat than BMI in the elderly (Crow et al., 2019). Of note is that even after adjusting for BMI, frailer women but not men also showed higher levels of sCD14, mostly because of an increase in this marker’s level in the preceding 20 years. sCD14 is a marker of monocyte activation since it is released from monocytes after stimulation with TLR ligands (Shive et al., 2015), in particular lipopolysaccharide (LPS). sCD14 also binds to LPS and it has been suggested that sCD14 is necessary for an adequate response to LPS in platelets (Damien et al., 2015) and can enhance sensitivity for LPS in endothelial cells (Lloyd-Jones et al., 2008), suggesting that sCD14 is important in the defense against bacterial infections. Its increase in concentration over time in frailer women is in line with previous reports showing higher monocyte numbers in frailer women (Leng et al., 2009; Samson et al., 2020).

While most associations between inflammatory marker levels and frailty were positive, as expected when low-grade inflammation occurs more frequently in frail participants, in women we also found a negative association of sCD40L levels with frailty, although only when not adjusted for BMI. sCD40L, which is released mainly by activated platelets (Danese, 2003; Nagasawa et al., 2005), is a soluble membrane glycoprotein involved in B cell responses (Jenabian et al., 2014) and in macrophage and monocyte signaling (Suttles & Stout, 2009). It thus plays an important role in the communication between innate and adaptive immune responses. sCD40L is commonly seen as a pro-inflammatory molecule (Aloui et al., 2016), but it might be immunosuppressive as was suggested in HIV infection and in cancer by inducing expansions of regulatory T cell populations (Huang et al., 2012; Jenabian et al., 2014), and suppressing T cell proliferation (Huang et al., 2012), suggesting that lower levels of sCD40L might indicate failing regulation with higher low-grade inflammation as a result.

### 3.3 Lung function and physical strength

Higher overall CRP levels are often associated with worse outcomes in specific health-related parameters, such as lung function (Ahmadi-Abhari et al., 2014) and handgrip strength (Cesari et al., 2014; Smith et al., 2019; Tuttle et al., 2020). We largely confirmed these results in our study, showing that inflammatory marker levels are persistently higher during 20 years in people with reduced lung function and handgrip strength, which also could indicate that low-grade inflammation in the elderly is indeed ‘chronic.’ In addition, only in women, higher levels of sCD40L and CXCL11/I-TAC were found to be associated with better lung function (by means of FEV1 and FVC), which we believe is the first time these markers were related to lung function in a community-dwelling population. While sCD40L was shown in previous studies to possibly have anti-inflammatory roles as discussed above, CXCL11/I-TAC was not, and we would have expected a negative correlation of this marker with lung function because of its pro-inflammatory effect in the IFNγ pathway. Such a negative association was seen in cross-sectionally performed studies in sarcoidosis (Arger et al., 2019) and other lung diseases (Kameda et al., 2020), showing higher levels of CXCL11/I-TAC with lower lung function, which is in contrast with our findings. Further studies are needed to confirm our findings in a community-dwelling population and to explain why in our study this relationship was found only in women.

Interestingly, in men we also found a negative correlation of handgrip strength with concentrations of IL-10, a cytokine commonly seen as anti-inflammatory. A similar relationship of higher muscle strength (knee extension strength) and lower IL-10 values was also described in a previous study (Calvani et al., 2017), although in most studies no associations between IL-10 and grip strength were detected (Cesari et al., 2004; Goldeck et al., 2016). Our results also show that IL-10 correlates strongly with several pro-inflammatory markers such as IL-6, which is in line with previous studies (Calvani et al., 2017; Hsu et al., 2019). Further research is needed to confirm the results and to investigate whether these elevated IL-10 levels could be a compensatory response to low-grade inflammation.

### 3.4 Strengths and limitations

Our longitudinal analysis of a well described cohort is a major strength of this study. To the best of our knowledge, our study is unique as an extensive inflammatory panel was repeatedly measured at 5-year intervals in the same individuals over a period of 20 years. Since variation within-individuals was much smaller than the variation between individuals, we had more power to detect changes of protein levels with advancing age than in cross-sectional studies, even when these have larger sample sizes. Furthermore, while sample size for cross-sectional analyses of a single analyte was limited compared to large cohort studies, our study made use of a large panel of inflammatory markers, thus giving a more comprehensive insight in inflammation than studies that measured only a few inflammatory markers. Also, since inflammatory marker levels can vary considerably between studies due to methodological differences, an advantage of our study was that all inflammatory markers were measured with the same assay and that all samples of the same individual were measured in the same plate. Of note is that we did not have a frailty index measurement at baseline, which is a limitation in our study of the relationship between levels of inflammatory markers and becoming frail. Thus, while we assumed that the participants who we selected to be healthy at baseline had a low frailty index score, this selection was based on a relatively simple health score rather than the frailty index score that we used at study endpoint. Other limitations are that the study population could be less representative for the general population due to, for instance, selective dropout of participants commonly seen in longitudinal cohorts, although the response rate in the Doetinchem cohort study is good, generally above 70% (Picavet et al., 2017). Furthermore, the prolonged storage of the plasma samples could affect the sample quality. However, the samples were consistently stored in low temperature and the trajectories were remarkably stable within individuals, indicating that protein degradation was probably of minor influence.

### 3.5 Concluding remarks

In conclusion, our exploratory study gives unique insight into longitudinal changes in the immune profile of aging men and women over a period of about 20 years, and revealed multiple associations of immune markers with clinically relevant outcomes at study endpoint, such as frailty, handgrip strength and lung function. More specifically, the findings implicate an important role for the IL-6 pathway due to elevated markers such as CRP and, only in women, monocyte activation due to the involvement of sCD14. As BMI and waist circumference are related to elevations of immune markers in the IL-6 pathway, chronic inflammation might be an important mediator of the relationship between BMI and frailty.

## 4 Methods

The Doetinchem cohort study (DCS) is a population-based longitudinal study of 7769 participants that have been followed since 1987 (Picavet et al., 2017; Verschuren et al., 2008). Every five years plasma samples were taken and data regarding the participants’ life style and health were collected (1). For the present study, a subgroup of 144 people aged 65-75 years was selected from the DCS, stratified by sex and with equal numbers of healthiest, intermediate and frailest participants according to the classification used in (Samson et al., 2019). In brief, the healthiest/frailest were defined as those with the 15% lowest/highest frailty index score (see below) among their age- and sex-matched peers. The individuals selected were participating in the study at least until 2016, and had participated in at least four out of the five investigated measurement rounds. The measurements of the present study were performed as an extension of the ones in the entire DCS, with the sample size being restricted by budgetary and logistic constraints.

### 4.1 Frailty index

Details on the frailty index score used in this study can be found elsewhere (Samson et al., 2019). Its definition was based on previous studies (Collerton et al., 2012; Mitnitski et al., 2001) and in our implementation it consists of 35 potential age- and health-related deficits, covering a broad range of ‘domains,’ including cognitive, physical, and psychological functioning. Difference of the score used in this study compared to the one previously used and validated in the DCS (Samson et al., 2019) is that the latest available data was included, and that the deficit regarding overweight was left out of the score, so that we could better investigate the influence of overweight on our results. Details and cutoff values for all of the 35 deficits are shown in Table S2. All deficits are bound between zero (best possible score) and one (worst possible), and the frailty index is the average score per individual. The frailty index was calculated in the two latest available DCS assessment rounds (rounds 5 and 6). Frailty categories (see DCS subgroup selection above) were only used for visualization in some figures; for statistical analyses, the continuous frailty index score was used.

### 4.2 Healthy aging index

The healthy aging index was measured at study baseline and was used to select a subgroup of participants that was ‘healthy’ at the start of follow-up. The score is based on health deficits defined by systolic blood pressure, plasma glucose levels, creatinine levels, lung function (forced vital capacity), and general cognitive function (Nooyens et al., 2011), as described elsewhere (Dieteren et al., 2020). Except for plasma glucose levels, the deficits were comparable to some of those included in the frailty index score. Every item has similar weight in the score, and the score ranges from 0 to 10, with 10 being the best and 0 the worst possible outcome (with all deficits present).

### 4.3 Measurements of inflammatory markers

Plasma samples were collected repeatedly within the same individuals over approximately 20 years between 1991-2017 (1). The samples had been stored at −80°C and thawed shortly before measuring a large panel of inflammatory markers, consisting of the following cytokines, chemokines and soluble receptors: C-C Motif Chemokine Ligand (CCL) 1/I-309, CCL2/MCP-1, CCL5/RANTES, CCL11/Eotaxin, CCL27/C-TACK, C-X-C Motif Chemokine Ligand (CXCL) 9 /MIG, CXCL10/IP-10, CXCL11/I-TAC, Interleukin (IL)-1β, IL-10, IL-6, soluble CD40 ligand (sCD40L), soluble CD14 (sCD14), soluble IL-2 receptor (sIL-2R), soluble IL-6 receptor (sIL-6R), soluble glycoprotein 130 (sGP130), Complement 5a (C5a), Brain-derived neurotrophic factor (BDNF), and P selectin. Concentrations of the markers were measured using a validated bead-based multiplex immunoassay (Flexmap 3D®, Luminex) at the Multiplex Core Facility lab, University Medical Center Utrecht, The Netherlands. All detection antibodies were coupled to magnetic beads and were tested for specificity. Samples were thawed at room temperature just before the measurement. To minimize assay variation within individuals, all samples from the same participant (n=6) were measured on the same plate.

Measurement of samples took place in 2018, in three independent batches. Levels of C-reactive protein (CRP) in plasma had been measured previously in DCS round 2,3,4, and 5 (Hulsegge et al., 2016). Apart from sIL-2R and IL-1β, the proportion of samples in which concentrations were below the limit of quantification (LOQ) was less than 20% for all inflammatory markers. Concentrations below the LOQ were imputed with a random value below the LOQ based on maximum likelihood estimation (Lubin et al., 2004). sIL-2R and IL-1β were excluded from analysis since more than 65% of the samples were below the LOQ. For IL-6 and IL-10, 85 samples (often in the same participants) were excluded due to possible non-specific binding to beads in the IL-10 or IL-6 region, which reduced the sample size for these cytokines to n=129 out of the total n=144 individuals (n=65 men, n=64 women). The sample size of sGP130 was reduced (n=67 men, n=66 women) since sGP130 was not measured in the first batch of cytokine measurements.

### 4.4 Anthropometric measurements

Body mass index was calculated as body mass in kilograms divided by the square of the participants’ height in meters. A tape measure was used to measure waist circumference, which was done with the participant in an upright standing position and holding the tape measure horizontal around the waist, at the midpoint between the lower rib and the iliac crest.

### 4.5 Handgrip strength

Participants applied as much force as possible with their dominant hand to a hydraulic dynamometer (Jamar), with their elbow in a 90 degree angle. The highest applied force out of three separate attempts was used.

### 4.6 Lung function

At study endpoint, a heated pneumotachometer (E Jaeger, Wurzburg, Germany) was used to measure forced expiratory volume in one second (FEV1) and forced vital capacity (FVC) as previously explained in detail (van Oostrom et al., 2018). Spirometric measurements were done and evaluated according to the American Thoracic Society and European Respiratory Society guidelines (Enright et al., 1991). FEV1 and FVC values were transformed to the percentage of predicted values, with predictions based on the Global Lung Initiative’s (2012) reference values (Quanjer et al., 2012) specific for age, length, sex, and ethnicity, using the R macro provided on https://www.ers-education.org/guidelines/global-lung-function-initiative/spirometry-tools.aspx.

### 4.7 Statistical analysis

#### 4.7.1 Longitudinal protein trajectory estimators

To estimate the “inflammatory burden over time” for each individual and each inflammatory marker the area under the concentration versus time curve (AUC) was calculated and was divided by the total time of follow-up to adjust for possible differences in follow-up time. The AUC values of every inflammatory marker were subsequently log transformed and mean-centered per measurement batch to adjust for possible confounding batch effects. To estimate the changes in inflammation over time, we also calculated the within-individual proportional increase or decrease per year. To do this, we used a slope of log-transformed concentration over time per individual, calculated in a median based linear model (Komsta, 2019), as an estimator for rate of change over time. A median based linear model is much less sensitive to outliers or to deviations from a normal distribution than ordinary linear regression models (Theil, 1950; Wilcox, 1998).

#### 4.7.2 Association studies

All the results were adjusted for multiple testing by controlling the false discovery rate (Benjamini & Hochberg, 1995) separately for each analysis. A nominal false discovery rate of at most 15% (meaning, roughly speaking, an average of 1.5 false discoveries per 10 discoveries presented as such) was tolerated in view of the number of tests carried out. For testing associations, we used the permutation versions of the Spearman and Wilcoxon-Mann-Whitney tests blocked for certain background or confounding variables (e.g. age) as implemented in the R package coin (Hothorn et al., 2008) with p-values estimated by simulation. For an explanation of the concept of blocking in experimental designs, see (Krzywinski & Altman, 2014). Details of all association studies are shown in Supplementary tables.

To investigate whether changes in plasma concentrations of inflammatory markers change with age, we carried out Spearman tests between each marker’s concentrations and age blocking by participant. Thus, we focused on within-individual increases in the marker’s concentration during the 20 year of follow-up. Should a marker tend to increase or decrease with age, the Spearman correlation between the marker’s concentration and an participant’s age should tend to exhibit consistently large or consistently small values across the cohort.

Single associations between an inflammatory marker’s trajectory (the AUC) and sex were tested with the Wilcoxon-Mann-Whitney test, blocking for age at last measurement (in two categories, 65-70 and 70-75 years) and BMI (three categories) to adjust for these variables.

Associations between inflammatory markers at the study’s endpoint were adjusted for age at last measurement and the immunoassay batch number (three batches). Longitudinal relationships between pairs of biomarkers were tested by investigating synchronous changes in biomarker levels. For every pair of markers, we investigated whether the moment of maximum increase in one marker coincided with that in another marker more often than what might be expected by chance, by carrying out one-tailed binomial tests postulating the ‘probability of success’ of ¼ under the null hypothesis of no association.

To address our research question whether chronic low-grade inflammation is related to frailty, we carried out several association studies, all separately for men and women. First, we tested associations, with Spearman’s tests, between the AUC of each marker’s inflammatory trajectory and the frailty index score, adjusted for age at last measurement. Analogous associations were tested using BMI as additional blocking variable, to get an idea about the importance of BMI in these relationships. Then, we investigated the association between the individual’s slope of the inflammatory protein trajectories and frailty and at endpoint, using Spearman’s tests blocked by the baseline level of the marker (in tertiles) and BMI. Next, the relation was investigated between every immune marker trajectory and change in frailty index score from round 5 to round 6, adjusting the associations for age at last measurement and for the initial frailty index score at round 5 (using tertiles for blocking). The latter was done to adjust for a possible regression to the mean effect and to give more weight to smaller increases in frailty index of people that already had a high index in the beginning.

The tested relationships of the inflammatory markers with lung function parameters were also adjusted for smoking with two blocking categories: non-smokers and former smokers. The relationships with handgrip strength were adjusted for age and were repeated with adjustments for BMI.

#### 4.7.3 Prediction analysis

We used a random forest prediction algorithm (Liaw & Wiener, 2002) to investigate whether frailty could be predicted using several dependent variables, namely the AUCs of all the inflammatory markers, age at endpoint, and BMI. The overall performance of the prediction analysis was assessed in terms of the percentage explained variance. The importance of the independent variables to predict frailty was assessed by ranking their ‘importance.’ This variable importance was quantified in terms of the percentage increase in mean-squared error (MSE) when the effect of that variable was removed.

All statistical analyses were performed with R version 3.6.2 (R Core Team, 2019), with several general packages for data processing (Wickham et al., 2020; Wickham & Henry, 2020) and for visualization (Pedersen, 2019; Wickham, 2016; Wilke, 2019). Correlations between inflammatory markers were visualized using the corrplot package (Wei & Simko, 2017).

## Supporting information

Supplementary correlation tables

## 5 Acknowledgments

We would like to thank the DCS respondents for their participation in the study and the epidemiologists and fieldworkers of the Municipal Health Service in Doetinchem for their contribution to the data collection. In addition, we would like to thank Petra Vissink for her help with the blood sample collection. Figure 1 and Figure 7a were created using BioRender.com.

## 6 Competing interests

None to declare.

## Supplementary information

**Table S1:** Summary characteristics of the cytokines and chemokines in the cohort

| Protein | Baseline concentration | Endpoint concentration | AUC |
| --- | --- | --- | --- |
| **C-reactive protein** |  |  |  |
| CRP | 1.03(0.93 – 1.14) * 10<sup>6</sup> | 1.22(1.11 – 1.34) * 10<sup>6</sup> | 1.56(1.43 – 1.70) * 10<sup>6</sup> |
| **CC chemokines** |  |  |  |
| CCL1/I-309 | 3.30(3.17 – 3.43) | 3.34(3.22 – 3.46) | 3.38(3.27 – 3.50) |
| CCL2/MCP-1 | 1.03(1.00 – 1.06) * 10<sup>2</sup> | 1.07(1.04 – 1.10) * 10<sup>2</sup> | 1.08(1.06 – 1.11) * 10<sup>2</sup> |
| CCL5/RANTES | 8.83(8.42 – 9.26) * 10<sup>4</sup> | 7.60(7.27 – 7.95) * 10<sup>4</sup> | 8.00(7.73 – 8.27) * 10<sup>4</sup> |
| CCL11/Eotaxin | 4.76(4.64 – 4.89) | 4.97(4.84 – 5.11) | 5.08(4.97 – 5.19) |
| CCL27/C-TACK | 9.34(8.97 – 9.74) * 10<sup>2</sup> | 1.03(0.99 – 1.08) * 10<sup>3</sup> | 1.08(1.05 – 1.12) * 10<sup>3</sup> |
| **CXC chemokines** |  |  |  |
| CXCL9/MIG | 3.08(2.95 – 3.23) | 2.98(2.85 – 3.11) | 3.15(3.05 – 3.26) |
| CXCL10/IP-10 | 2.37(2.29 – 2.46) * 10<sup>2</sup> | 2.83(2.73 – 2.93) * 10<sup>2</sup> | 3.01(2.92 – 3.10) * 10<sup>2</sup> |
| CXCL11/I-TAC | 6.07(5.64 – 6.53) | 6.19(5.76 – 6.65) | 6.98(6.59 – 7.39) |
| **Interleukins** |  |  |  |
| IL-6 | 2.43(2.23 – 2.65) | 2.31(2.13 – 2.51) | 2.61(2.45 – 2.78) |
| IL-10 | 0.71(0.66 – 0.77) | 0.66(0.62 – 0.72) | 0.75(0.71 – 0.80) |
| **Soluble receptors** |  |  |  |
| sCD14 | 2.15(2.10 – 2.21) * 10<sup>6</sup> | 2.21(2.15 – 2.27) * 10<sup>6</sup> | 2.27(2.22 – 2.32) * 10<sup>6</sup> |
| sCD40L | 4.14(3.90 – 4.40) * 10<sup>2</sup> | 3.69(3.48 – 3.91) * 10<sup>2</sup> | 4.05(3.87 – 4.24) * 10<sup>2</sup> |
| sIL-6R | 2.27(2.22 – 2.32) * 10<sup>4</sup> | 2.35(2.30 – 2.41) * 10<sup>4</sup> | 2.37(2.33 – 2.42) * 10<sup>4</sup> |
| **Other** |  |  |  |
| P selectin | 7.95(7.43 – 8.50) * 10<sup>4</sup> | 6.80(6.40 – 7.22) * 10<sup>4</sup> | 8.04(7.73 – 8.36) * 10<sup>4</sup> |
| sGP130 | 2.19(2.16 – 2.22) * 10<sup>4</sup> | 2.23(2.20 – 2.26) * 10<sup>4</sup> | 2.25(2.23 – 2.27) * 10<sup>4</sup> |
| C5a | 4.65(4.38 – 4.93) * 10<sup>3</sup> | 4.90(4.62 – 5.18) * 10<sup>3</sup> | 5.06(4.79 – 5.34) * 10<sup>3</sup> |
| BDNF | 1.85(1.79 – 1.91) * 10<sup>4</sup> | 1.69(1.63 – 1.77) * 10<sup>4</sup> | 1.75(1.69 – 1.81) * 10<sup>4</sup> |

*Note:*
Concentrations ( $pg * mL^{-1}$ ) and AUC values ( $years * pg * mL^{-1}$ ) are geometric mean values with 95% confidence intervals.

**Table S2:** Frailty index components

| No | Frailty index component | Description | Value: 0 | Value: 0.5 | Value: 1 |
| --- | --- | --- | --- | --- | --- |
| 1 | RR | High (systolic) blood pressure | $RR < 160$ | | $RR \geq 160$ |

Table S2: Frailty index components (*continued*)
| No | Frailty index component | Description | Value: 0 | Value: 0.5 | Value: 1 |
| --- | --- | --- | --- | --- | --- |
| 2 | Cardiologic Disease | One or more of the following conditions: myocardial infarction, bypass-surgery, balloon dilatation, cardiac catheterization, pacemaker implantation, large blood vessel surgery, hospitalization due to cardiac failure | No |  | One or more prevalent |
| 3 | Diabetes | Prevalence of Diabetes | No |  | Yes |
| 4 | Hearing | Inability to maintain a conversation in a group of 3 or more people due to hearing impairment (with hearing aid if needed) | Yes or with some effort |  | No or with great effort |
| 5 | Malignancy | (History of) any form of malignancy | No |  | Yes |
| 6 | Joint Inflammation | Chronic joint inflammation in the past year | No |  | Yes |
| 7 | Osteoporosis | Osteoporosis diagnosed by a medical doctor, in the past year, | No |  | Yes |
| 8 | Lower Back Pain | Severe lower back complaints in the past year (including lumbar herniated nucleus pulposus) | No |  | Yes |
| 9 | CVA | (History of) stroke | No |  | Yes |
| 10 | Migraine | Migraine prevalence in the past year | No |  | Yes |

Table S2: Frailty index components (*continued*)
| No | Frailty index component | Description | Value: 0 | Value: 0.5 | Value: 1 |
| --- | --- | --- | --- | --- | --- |
| 11 | Neurologic Disease | One or more of the following neurological diseases, diagnosed by a medical doctor: m. Parkinson, Multiple Sclerosis, epilepsy | No |  | Yes |
| 12 | Asthma | Asthma diagnosed by a doctor and one or more asthma attacks in the past year | No |  | Yes |
| 13 | Spirometry Ratio | Poor lung function quantified by spirometry measurements: first second of forced expiration divided by the forced vital capacity (FEV1/FVC) | $FEV1/FVC > 0.70$ | | $FEV1/FVC \leq 0.70$ |
| 14 | Digestive Tract | Severe bowel disorders in the past year, diagnosed by a medical doctor | No |  | Yes |
| 15 | Vertigo | Vertigo with falling the past 12 months | No vertigo | Some vertigo | Severe vertigo |
| 16 | Pain | Limited in daily activities due to pain | No limitation | Some limitation | Severe limitation |
| 17 | Incontinence | Unintentional urine incontinence past 12 months | No |  | Yes |
| 18 | Ankle Brachial Index | Poor ankle brachial index (ABI, the ratio of the systolic blood pressure in the ankle to the blood pressure in the arm) | $ABI > 0.9$ | | $ABI \leq 0.9$ |

Table S2: Frailty index components (*continued*)
| No | Frailty index component | Description | Value: 0 | Value: 0.5 | Value: 1 |
| --- | --- | --- | --- | --- | --- |
| 19 | Health Perception | (Subjective) perception of poor health | No or some impairment |  | Severe impairment |
| 20 | Eyesight | Bad eyesight perception, facial recognition within 4 meters (with glasses/ contact lenses if needed) | No |  | Yes |
| 21 | Renal Function | Poor renal function (estimated Glomerular Filtration Rate (eGFR), calculated with plasma creatinine concentrations) (Inker et al., 2012) | $eGFR \geq 60$ | | $eGFR < 60$ |
| 22 | Cognitive Speed | Poor cognitive speed. Z-scores corrected for measurements per person and for education level. Scores derived from the Stroop Color-Word Test and the Letter-Digit Substitution Test, as described previously (Nooyens et al., 2011) | Not belonging to 10% participants with lowest z-score within the Doetinchem cohort |  | Belonging to 10% participants with lowest z-score within the Doetinchem cohort |
| 23 | Cognitive Memory | Poor cognitive memory. Z-scores corrected for measurements per person and for education level. Scores derived from the Verbal Learning Test, as described previously (Nooyens et al., 2011) | Not belonging to 10% participants with lowest z-score within the Doetinchem cohort |  | Belonging to 10% participants with lowest z-score within the Doetinchem cohort |

Table S2: Frailty index components (*continued*)
| No | Frailty index component | Description | Value: 0 | Value: 0.5 | Value: 1 |
| --- | --- | --- | --- | --- | --- |
| 24 | Cognitive Flexibility | Poor cognitive flexibility. Z-scores corrected for measurements per person and education level. Scores derived from the Stroop Color-Word Test, as described previously (Nooyens et al., 2011) | Not belonging to 10% participants with lowest z-score within the Doetinchem cohort |  | Belonging to 10% participants with lowest z-score within the Doetinchem cohort |
| 25 | Physical Inactive | Not meeting the Dutch healthy exercise norm.(Kemper, 2000). In addition: belonging to the 25th lowest percentile of walking activity and the 10th percentile lowest low/medium/high intensive activities in the Doetinchem cohort | No |  | Yes |
| 26 | ADL | Limited in washing and dressing due to health | No limitation | Some limitation | Severe limitation |
| 27 | Household | Limited in daily activities (cooking, cleaning) due to health | No limitatlion | Some limitation | Severe limitation |
| 28 | Walking | Limited in in walking 100 meters | No limitatlion | Some limitation | Severe limitation |
| 29 | Lifting | Limited in lifting or carrying groceries due to health | No limitatlion | Some limitation | Severe limitation |

Table S2: Frailty index components (*continued*)
| No | Frailty index component | Description | Value: 0 | Value: 0.5 | Value: 1 |
| --- | --- | --- | --- | --- | --- |
| 30 | Walking Stairs | Limited in climbing stairs | No limitatlion | Some limitation | Severe limitation |
| 31 | Grip Strength | Poor grip strength (cutoff points as described previously) (Fried et al., 2001) |  |  |  |
| | | *Men, BMI* $\leq 24$ | $> 29$ | | $\leq 29$ |
| | | *Men, 24* $< BMI \leq 26$ | $> 30$ | | $\leq 30$ |
| | | *Men, 26* $< BMI \leq 28$ | $> 30$ | | $\leq 30$ |
| | | *Men, BMI* $> 28$ | $> 32$ | | $\leq 32$ |
| | | *Women, BMI* $\leq 24$ | $> 17$ | | $\leq 17$ |
| | | *Women, 23* $< BMI \leq 26$ | $> 17.3$ | | $\leq 17.3$ |
| | | *Women, 26* $< BMI \leq 29$ | $> 18$ | | $\leq 18$ |
| | | *Women, BMI* $> 29$ | $> 21$ | | $\leq 21$ |
| 32 | Depressed | Feeling depressed the past week | No limitatlion | Some limitation | Severe limitation |
| 33 | Happiness | Feeling unhappy the past 4 weeks | No limitatlion | Some limitation | Severe limitation |
| 34 | Mental Effort | Feeling as if every activity costs effort during the past week | No limitatlion | Some limitation | Severe limitation |
| 35 | Getting Going | Not being able to get going the past week | No limitatlion | Some limitation | Severe limitation |

**Figure S1:**
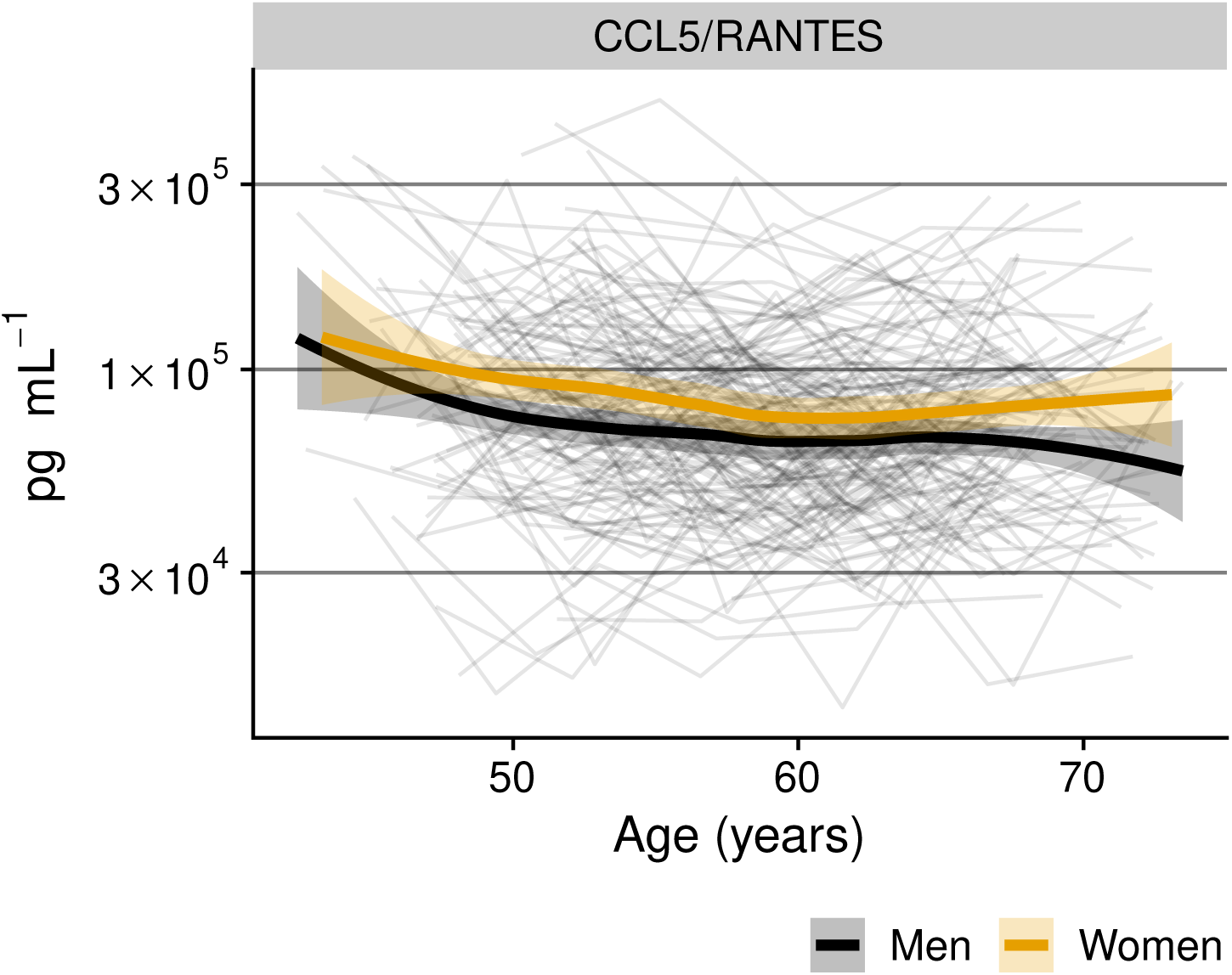
The concentration of CCL5/RANTES over 20 years in men and women (showing that this is continuously higher in women).

**Figure S2:**
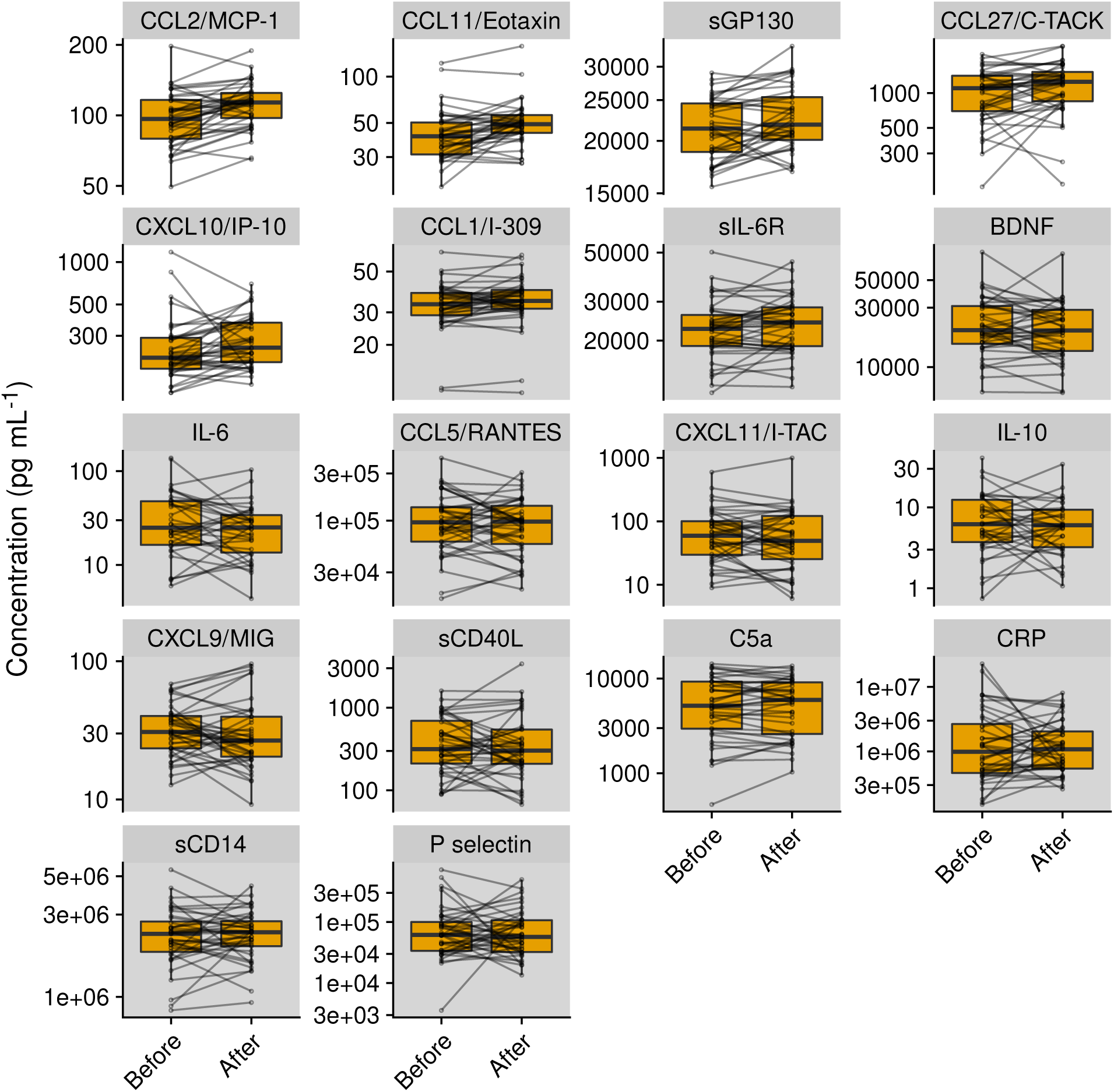
Menopause is related to changes in inflammatory marker profile. Inflammatory marker concentrations are shown shortly before and shortly after menopause (average difference: 5.3 years) in women of whom data at both timepoints are available (n=40/70). Tiles of biomarkers in which an association was found with menopause are shown in a white background, others in a grey background.

**Figure S3:**
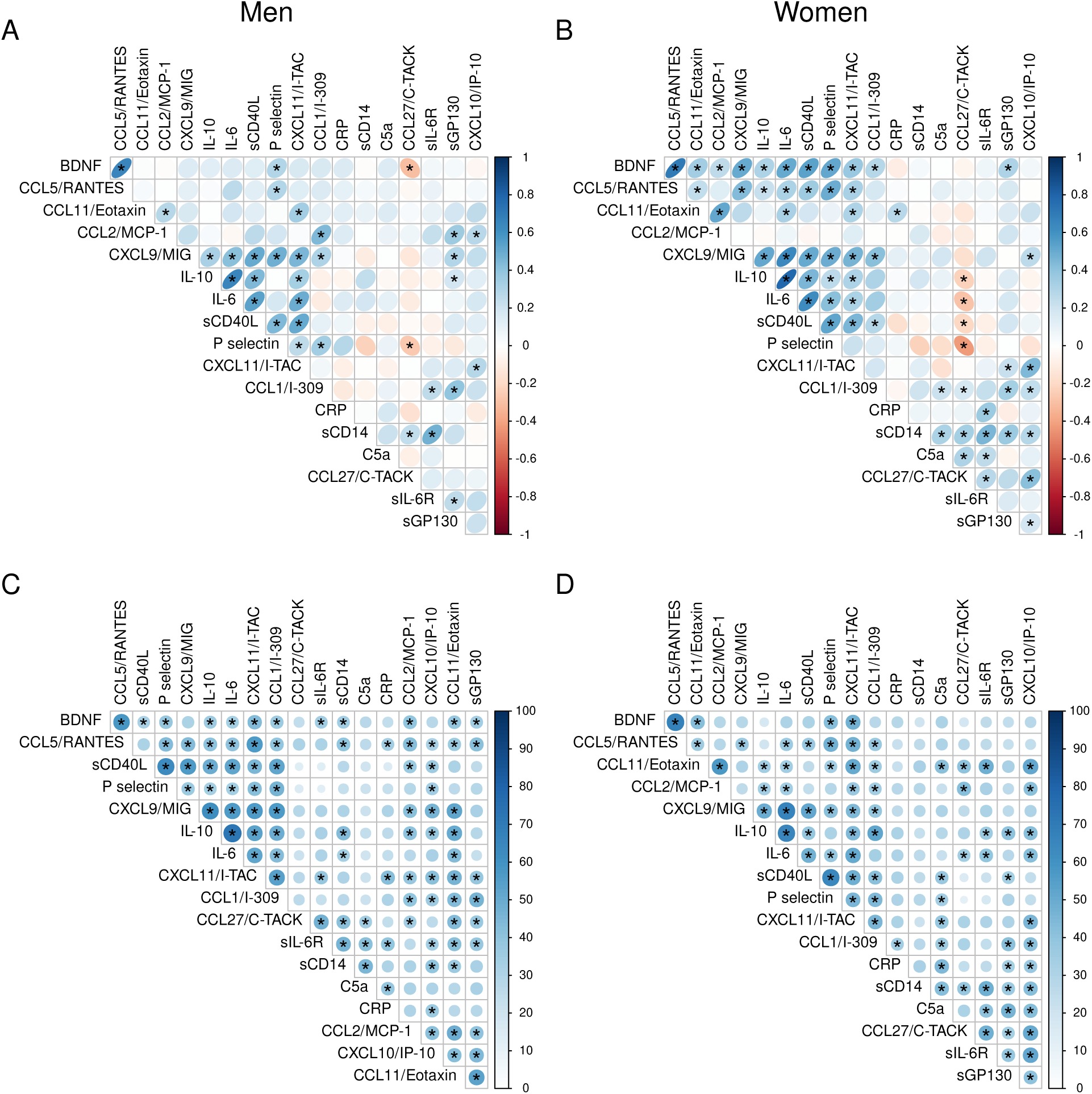
Relationships between inflammatory markers shown as (A,B) correlation between pairs of inflammatory markers at study endpoint and (C,D) similarity between pairs of inflammatory marker trajectories during about 20years of follow-up. In (A,B) the direction and strength of the association in is visualized with an oval shape and a color gradient. In (C,D) the blue gradient color and the size of the circles shows the percentage of participants of which a pair of biomarkers had the highest increase in concentration at the same moment in 20 years of follow-up. *= an association between two inflammatory markers, with false discovery rate being set at a maximum of 15%. n=71 women, n=73 men.

**Figure S4:**
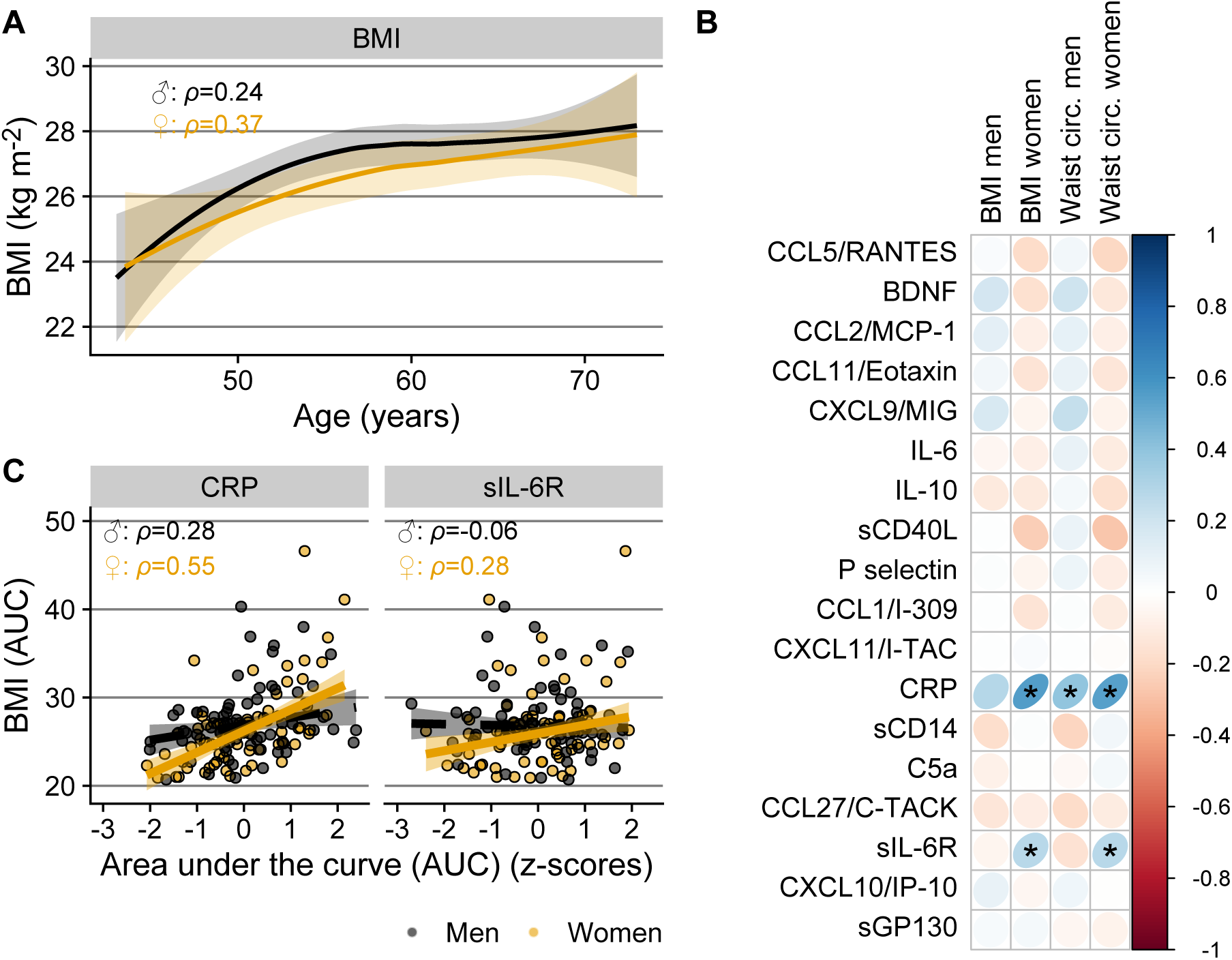
Relationship between body mass and inflammatory markers. (A) Local polynomial regression lines showing change in BMI over time in men and women. (B) Relationship of inflammatory marker levels with BMI and waist circumference of men (n=73) and women (n=71). Direction and strength of the associations in (B) are visualized with an oval shape and a color gradient. To capture the cumulative “exposure,” both BMI and inflammatory markers in (B) and (C) are expressed as area under the curve of levels/concentrations versus time, standardized to take into account different follow-up periods, and transformed into z-values. (C) BMI values (AUC) related to the AUC of CRP and sIL-6R. Trendlines in (C) are (robust) linear regression lines with 95% confidence interval.

## Notes

### Competing Interest Statement

The authors have declared no competing interest.

